# Classification of image category based on spatially distributed, transient high-frequency events

**DOI:** 10.64898/2026.05.15.725481

**Authors:** Alan Díaz, Idan Tal, Noah Markowitz, Shany Grossman, Elizabeth Espinal, Gelana Tostaeva, Charles Schroeder, Ashesh Mehta, Samuel Neymotin, Stephan Bickel

## Abstract

Transient high-frequency activity (HFA) in local field potentials exhibits substantial variability across trials and behavioral states, yet its functional role in sensory representation remains poorly understood. Here, we tested whether transient HFA events carry stimulus-related information through their temporal, spectral, and phase-dependent properties.

Using intracranial recordings from 21 human participants performing a visual localizer task, we extracted transient HFA events and quantified their features using information-theoretic and decoding analyses. Individual event features carried only modest information about stimulus identity, and response magnitude and morphology-related properties contributed minimally to decoding performance. Instead, decoding was dominated by temporal alignment and low-frequency phase. Critically, representing transient events as distributed low-frequency phase configurations across electrodes substantially improved cross-trial decoding performance, whereas disrupting distributed phase structure eliminated this decoding advantage.

These findings indicate that stimulus-related structure in transient HFA does not primarily arise from isolated local event properties, but instead emerges through distributed, phase-dependent network dynamics. More broadly, the results provide a framework for understanding how transient high-frequency neural activity contributes to sensory representations across distributed cortical networks.

## Introduction

Temporally structured neural activity is hypothesized to play a crucial role in interareal communication and the temporal modulation of excitability in the brain^1,2^. High-frequency activity (HFA) exhibits substantial temporal variability across trials and behavioral states, yet analyses of HFA rely on averaging signals across time and trials, emphasizing sustained activity over transient dynamics^3^. However, growing evidence suggests that high-frequency neural activity often occurs in brief, transient events, and that the dynamics within and across these events may carry more detailed information than averaged power estimates^4–9^.

Although adding noise to a continuous sinusoidal signal can produce apparent transient event structure with varying spectral properties across trials^3^, recent studies have raised the possibility that at least some transient high-frequency events detected in human intracranial recordings may reflect fluctuations in broadband aperiodic activity rather than discrete oscillatory events^10^. Nevertheless, it remains unclear to what extent transient HFA observed during perceptual tasks reflects structured stimulus-related processing rather than non-specific variability. While most prior work has focused on averaged HFA, the role of transient events in representing stimulus-specific information in naturalistic perceptual settings remains underexplored. Addressing this gap is essential for understanding how neural signals encode complex sensory inputs.

Low-frequency activity, particularly in delta and theta ranges, is thought to provide temporally extended excitability windows that facilitate large-scale coordination across cortical networks^11,12^. HFA occurring within these phase-defined windows may therefore reflect not only local activation, but also the distributed network state in which that activation emerges.

One mechanism that may support such organization is the temporal coordination of neural activity by ongoing low-frequency dynamics. In these frameworks, the timing of transient high-frequency events relative to ongoing low-frequency phase may influence neural excitability, interareal coordination, the structure and separability of stimulus representations^11,12^. Transient HFA events may therefore be informative not only in isolation, but also in relation to the distributed low-frequency network state in which they occur. We therefore hypothesized that the low-frequency phase associated with transient HFA events, particularly when considered across distributed cortical regions, provides a key dimension for encoding and decoding visual stimuli.

Here, we test this hypothesis using intracranial recordings during a visual localizer task, combining HFA event-based feature extraction with information-theoretic and decoding analyses. We show that individual event features carry limited but reliable information about stimulus identity and that decoding performance is not driven by response magnitude or spectral morphology. Instead, stimulus-related information is concentrated in low-frequency phase and temporal alignment, and is most effectively captured when represented as distributed phase configurations across electrodes.

Consistent with prior work showing that both broadband spiking activity and coherent oscillatory synaptic potentials contribute to high-frequency signals in local field potentials (LFPs) across micro- and meso-scale networks^13–18^, our findings support a view of transient HFA as a multi-scale signal shaped by interactions between local activity and distributed network state. By linking transient high-frequency events to distributed phase structure, this study provides a framework for understanding how temporally precise and spatially distributed neural dynamics contribute to sensory processing.

## Results

Our objective was two-fold: to investigate (i) whether electrophysiological high-frequency activity (70-150 Hz) serves as a neural marker of visual processing in the human brain, and (ii) whether this activity encodes information relevant to image categorization. To this end, we analyzed intracranial electroencephalography (iEEG) recordings from 21 patients with medically refractory epilepsy performing a passive visual localizer task.

During their stay in the epilepsy monitoring unit at North Shore University Hospital (Manhasset, NY, USA), patients viewed images presented for 240 ms across 360 trials, followed by a jittered inter-trial interval averaging 750 ms (**Figure 1A**). Stimuli belonged to six categories: animals, patterns, people, places, tools, and words, with 60 unique images per category. To maintain engagement, participants responded to occasional image repetitions, although our primary focus was on neural activity during passive viewing.

**Figure 1.**
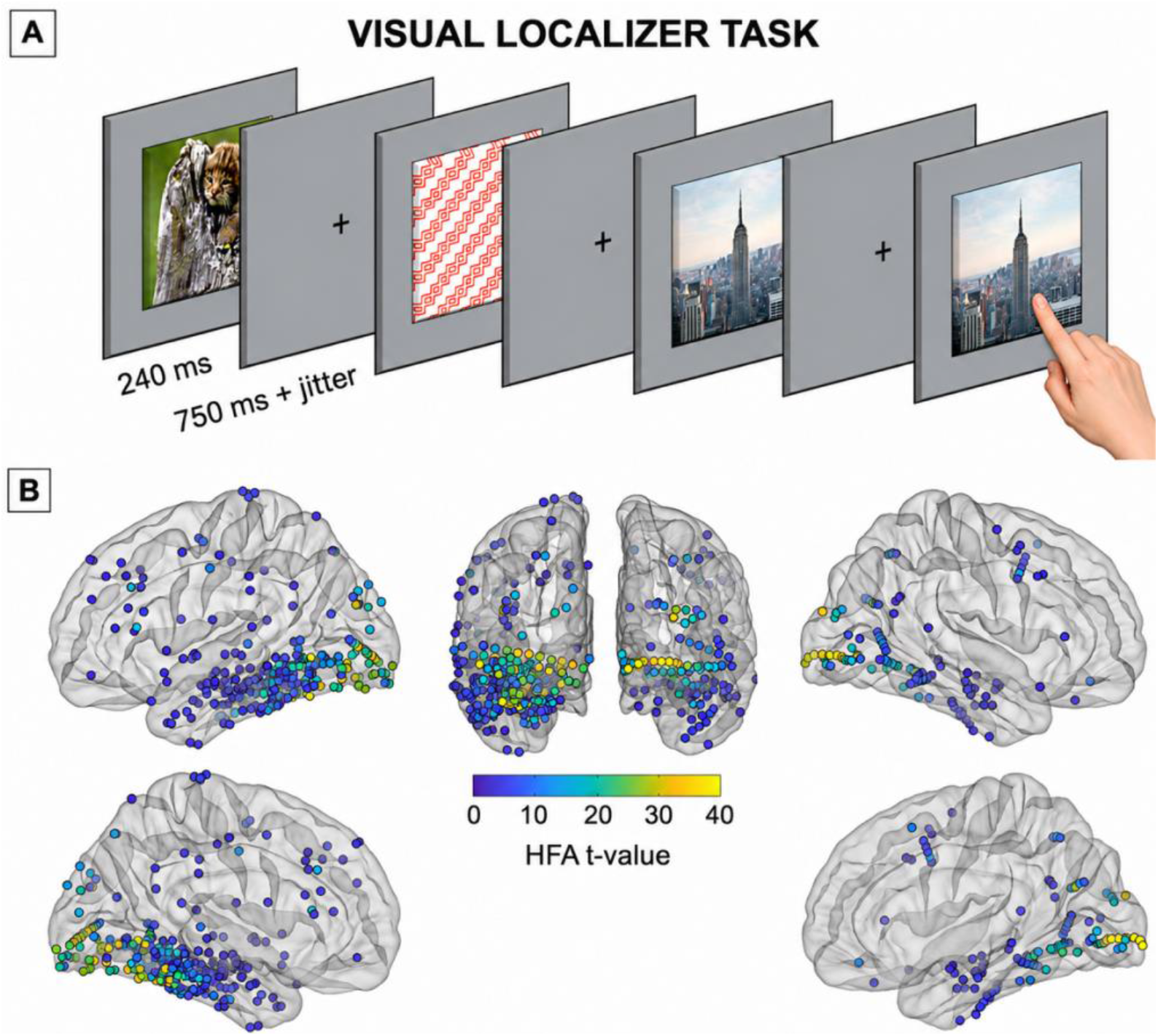
Visual task paradigm and distribution of stimulus-responsive HFA electrodes. **A**. Exemplar images are shown for the categories animals, patterns and places (left to right). Participants were prompted with an image for 240 ms followed by a jittered inter-trial period of 750 ms. Upon image repetition, participants pushed a button. **B**. Significant HFA (70-150 Hz) electrodes across all patients are shown on an average brain and color-coded based on their t-value vs. baseline. The highest t-values are found in occipital and ventral temporal regions but significant increase in HFA following the onset of image presentation was detected in a range of other temporal and frontal regions in both hemispheres.

We hypothesized that the brain automatically organizes visual stimuli into categorical representations upon presentation, and that such structure is reflected in high-frequency activity. Rather than assuming that the magnitude of the evoked response alone determines the informativeness of a given electrode, we considered the possibility that stimulus-related information may be encoded in more subtle aspects of the neural response.

### Identification of visually responsive electrodes

Because electrode placement varied across patients, we first identified electrodes exhibiting significant HFA increases following stimulus onset relative to baseline (t-test, p < 0.05). Across all subjects, 14.7% of electrodes (481/3270) showed significant responses and were termed *significant electrodes* for subsequent analyses.

As expected, the strongest responses were predominantly observed in occipital cortex, consistent with early visual processing. However, significant HFA increases were also detected across temporal and frontal regions (**Figure 1B**), indicating that visually driven activity extends beyond primary sensory areas.

### Response magnitude and category specificity

We next examined whether electrodes with the strongest evoked responses were also the most informative for distinguishing image categories. To this end, we compared response structure across electrodes with varying levels of stimulus-evoked HFA. **Figure 2** illustrates representative time-frequency responses from two electrodes with markedly different response magnitudes. The high-response electrode exhibited a robust, broadband increase in activity that was largely consistent across stimulus categories, reflecting a stereotyped evoked response. In contrast, the lower-response electrode displayed more heterogeneous and temporally structured activity that varied across categories.

**Figure 2.**
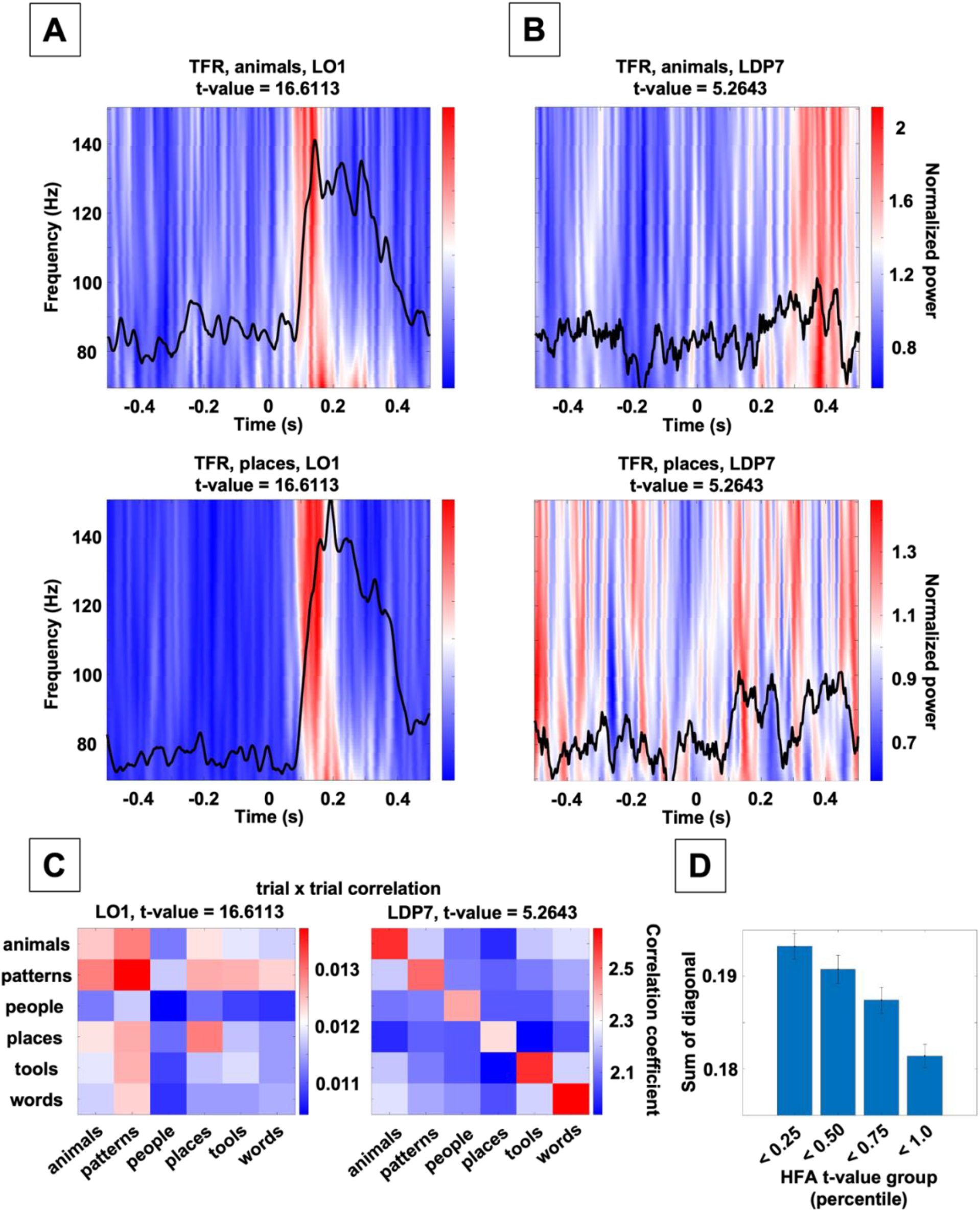
Time-frequency correlation structure across electrodes. **A-B**. Time-frequency representations for two example image categories from a high-response electrode (A) and a low-response electrode (B). Black traces show the evoked low-frequency signal (1-30 Hz). The t-value in each panel reflects the electrode’s overall response strength across all image categories. **C**. Trial-by-trial cross-correlation matrices of time-frequency maps within each image category. The high-response electrode shows more uniform correlation structure across trials, whereas the low-response electrode exhibits greater variability. **D**. Sum of diagonal elements of the correlation matrices across electrodes and subjects. Lower values indicate increased differentiation across trials, consistent with greater category specificity.

To quantify this observation, we computed trial-by-trial cross-correlations of time-frequency representations within each category. Electrodes with stronger responses exhibited higher within-category similarity, whereas electrodes with weaker responses showed more differentiated patterns across trials (**Figure 2C**). When electrodes were grouped by response magnitude, the average similarity within categories decreased as response strength increased (**Figure 2D**), indicating that strong evoked responses are associated with more stereotyped, less category-specific activity patterns. Importantly, this does not imply that weaker responses are intrinsically more informative. Rather, these findings indicate that response magnitude alone does not capture the dimensions of neural activity that differentiate stimulus categories and suggest that informative structure may reside in more complex aspects of the signal. This observation motivates a shift from analyzing response magnitude to explicitly quantifying the information carried by different components of HFA dynamics. In the following sections, we test this hypothesis using feature-based representations and information-theoretic and decoding analyses.

### Motivation for feature-based analysis of HFA dynamics

To move beyond response magnitude and capture the structure of HFA signals, we extracted transient high-frequency events using the Better Oscillation Detection Method (BOSC^19^) and quantified features describing their temporal, spectral, and state-dependent properties.

For each detected event, we extracted nine features: probability of occurrence, count, latency, center frequency, frequency span, number of cycles, duration, amplitude, and low-frequency phase (**Figure 3A**). Together, these features provide a multidimensional representation of transient HFA events, capturing not only their magnitude but also their timing, spectral content, and relationship to ongoing network state.

**Figure 3.**
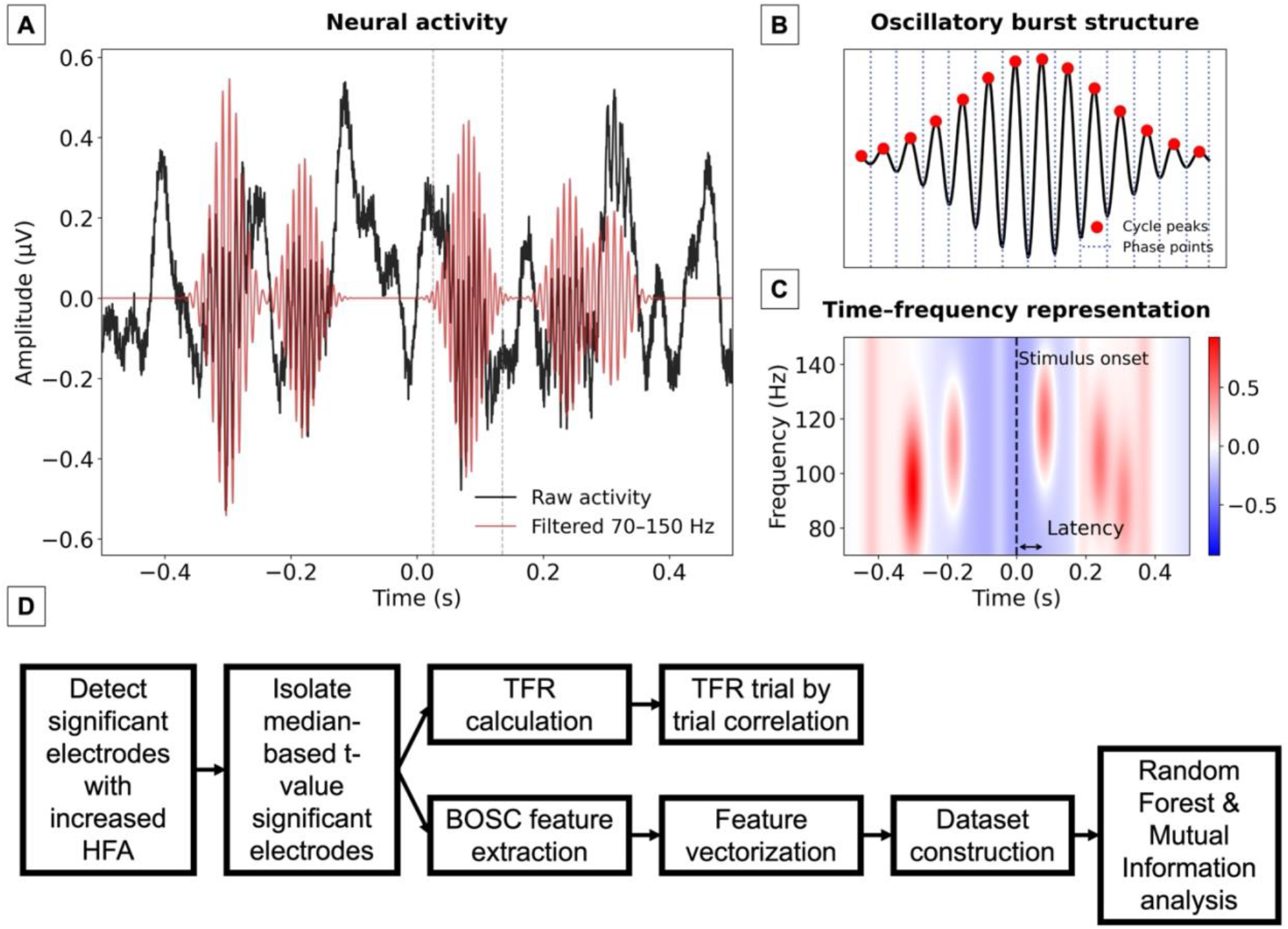
Feature-based representation of HFA dynamics. **A**. Example neural activity from a single electrode during visual stimulation, showing raw signal (black) and broadband high-frequency activity (70-150 Hz; red). **B**. Zoomed segment illustrating the transient structure of HFA. Individual cycles are identified by their peaks (red markers), and phase is defined relative to these transient components (blue dashed lines). Transient HFA event amplitude is quantified as the peak deviation from baseline within the event. **C**. Time-frequency representation of the same signal, showing transient increases in high-frequency power. Event latency is defined as the temporal offset of these events relative to stimulus onset. Together, these representations illustrate how multiple complementary features, capturing temporal, spectral, and dynamical properties of transient events, are extracted from HFA signals for subsequent analysis. **D**. Schematic summary of the event feature extraction framework. From HFA signals, multiple complementary features are quantified, including spectral morphology (e.g., cycle structure, amplitude, duration, center frequency, and frequency span), temporal properties (e.g., latency relative to stimulus onset), and phase-dependent dynamical structure. These extracted features provide a multidimensional representation of transient events and form the basis for subsequent decoding and information-theoretic analyses of neural population activity.

We constructed feature matrices by vectorizing these event-level representations across electrodes and trials, yielding labeled datasets in which each observation corresponded to a transient event associated with a specific stimulus category. Using these representations, we evaluated how different aspects of HFA dynamics contribute to stimulus-related information and image classification, using Random Forest models and permutation-based statistical evaluation (**Figure 3B**).

As a first step, we quantified the contribution of individual transient event features to stimulus identity, as suggested by the correlation analyses in **Figure 2**.

To isolate the contribution of individual transient HFA event features while preserving the spatial structure of the recordings, we constructed a feature representation in which each descriptor was evaluated independently across electrodes and trials. This formulation allows us to assess whether stimulus-related information is present at the level of single feature dimensions, and how this information varies across subjects and recording locations. We refer to this representation as Dataset 1.

Dataset 1 captures transient HFA event features at the level of individual trials and electrodes. For each subject, transient HFA events were detected independently at each electrode and aligned across the 360 image-presentation trials, yielding matrices with trials as rows and electrodes as columns. Each entry represents the value of a given feature at a specific electrode during that trial, with trials labeled by image category.

From each detected transient HFA event, we extracted nine descriptive features: event probability of occurrence, latency relative to stimulus onset, duration, event count, amplitude, center frequency, frequency span, event cycle count, and low-frequency phase. Accordingly, Dataset 1 consisted of nine feature-specific matrices, each containing 360 trials with electrode-wise feature values as columns.

Because the decoding task involved six image classes, the probability of correctly identifying the stimulus by chance is 1/6 (16.67%). In information-theoretic terms, this corresponds to a total stimulus uncertainty of log_2_(6) ≈ 2.58 bits prior to observing any neural response. We next quantify how much of this uncertainty is reduced by individual transient-event features using mutual information.

To quantify the stimulus-related information carried by each transient-event descriptor, we computed the mutual information (MI) between each feature representation X and stimulus identity Y across trials. For transient features that may be absent on some trials, the feature representation was defined as the joint variable

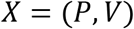

where P is a binary variable indicating event presence and V is the feature value conditional on event occurrence. Using the chain rule of mutual information, the total stimulus information carried by each feature was expressed as

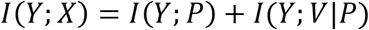

where Y denotes stimulus identity. This decomposition allowed us to separately quantify information conveyed by event occurrence and by the magnitude of the event feature itself.

To account for finite-sample bias, information estimates were evaluated relative to permutation-based null distributions generated by randomly shuffling stimulus labels across trials (200 permutations). ΔMI values correspond to the difference between the observed mutual information and the mean mutual information obtained from the shuffled-label distributions.

Across subjects, individual transient HFA event features carried modest but reliable stimulus information. Raw mutual information values were typically on the order of ∼0.10-0.14 bits per feature (**Figure 4B**), whereas permutation-corrected values (ΔMI) generally ranged from ∼0.03 - 0.08 bits per feature. While small relative to the total stimulus entropy (2.585 bits), these values were consistently above permutation-based null distributions, indicating the presence of structured, stimulus-related signal at the level of individual descriptors.

**Figure 4.**
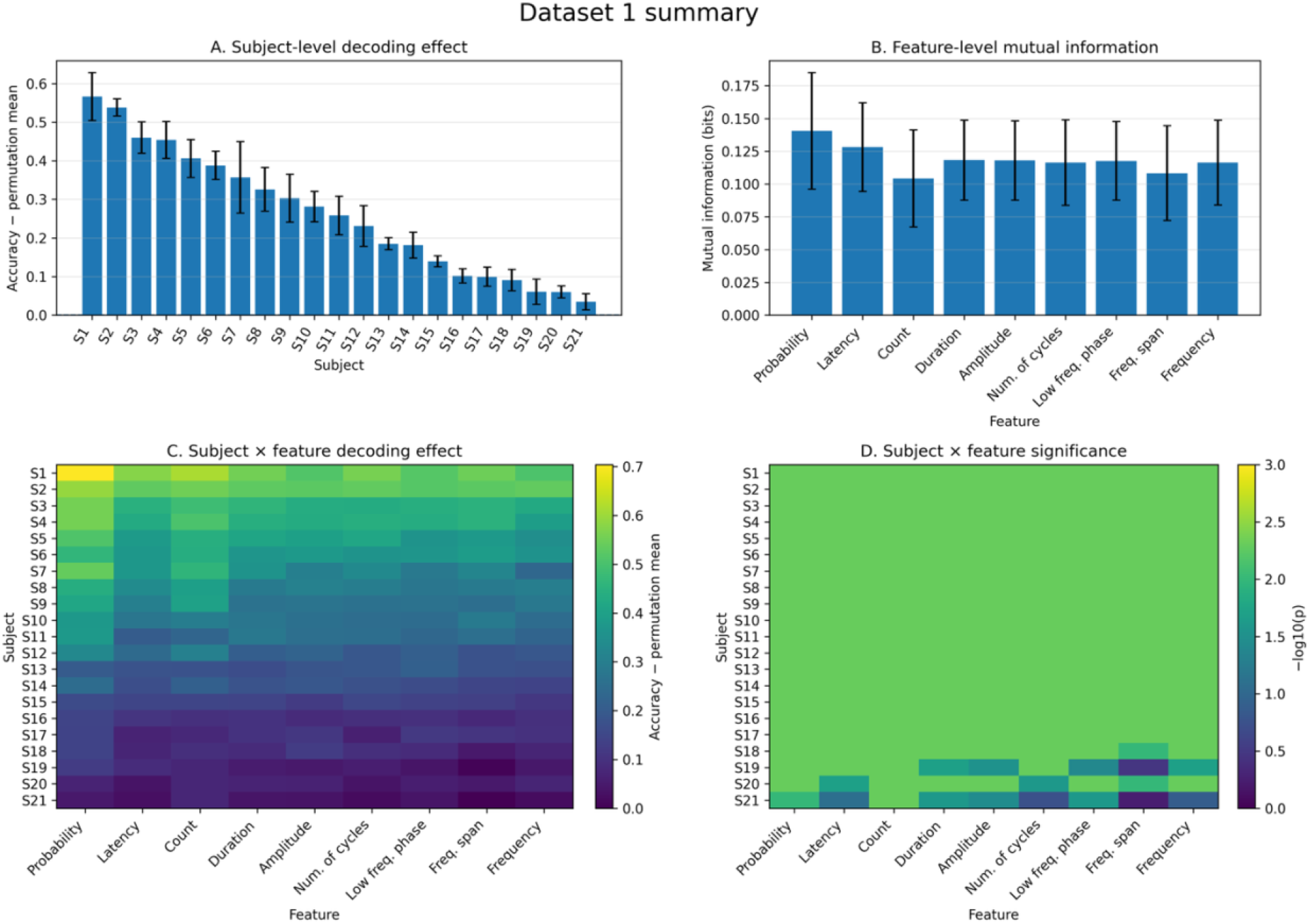
Dataset 1 summary of feature-level information and decoding performance. **A**. Subject-level decoding effect, defined as classification accuracy minus permutation-based mean accuracy, averaged across features. Subjects are ordered by mean decoding performance, revealing substantial variability across individuals. **B**. Mutual information between each transient HFA event feature and stimulus identity, averaged across subjects. All features carry measurable information, with probability, latency, and count showing slightly higher values, although differences across features are modest. Error bars indicate standard deviation across subjects. **C**. Heatmap of decoding effect size for each subject-feature pair. Strong horizontal structure indicates that subject-level variability dominates over feature-level differences, with higher-performing subjects showing elevated decoding across features. **D**. Corresponding heatmap of statistical significance, expressed as −log10(p). A value of 1.30 corresponds to p = 0.05. Most subjects exhibit significant decoding across features, whereas lower-performing subjects show weaker and less consistent effects.

At the feature level, transient HFA event probability showed the highest mutual information with stimulus identity, followed by latency. Most remaining features, including duration, amplitude, number of cycles, low-frequency phase, and center frequency, showed broadly comparable values, whereas event count and frequency span were slightly lower. When corrected against permutation-based estimates, the same general ordering was preserved, with probability, latency, and count showing the largest ΔMI values. Thus, stimulus-related information was not confined to a single transient-event descriptor but was distributed across multiple dimensions of transient HFA dynamics.

Taken together, these results indicate that stimulus-related information is distributed across features, with no single descriptor providing sufficient information for reliable classification. This supports the interpretation from **Figure 2** that category-specific structure is not captured by response magnitude alone.

A comparison of mutual information values across subjects and features further clarifies this structure (**Figure 4**). Transient HFA event probability, latency, and event count consistently showed the highest information values, while other features clustered at slightly lower but comparable levels. Notably, information values varied substantially across subjects for all features, indicating that differences in electrode coverage and anatomical sampling strongly constrain the amount of stimulus-related information that can be captured. Consistent with the decoding results, this suggests that variability in performance is driven primarily by where signals are recorded rather than by the intrinsic informativeness of individual features.

This demonstrates that stimulus information is reliably decodable from transient HFA event features at the electrode level, despite the relatively small contribution of any individual descriptor. The distributed nature of this information, combined with strong subject-level variability, indicates that category-relevant structure depends more on spatial sampling than on any single feature dimension, motivating a targeted analysis of how features jointly contribute to decoding performance.

These results raise a critical question: if stimulus-related information is distributed across features but individually weak, do all transient HFA event features contribute equally to decoding performance, or is information concentrated in specific dimensions of the response? To address this, we grouped features according to their temporal, spectral, and state-dependent properties and evaluated their contributions to classification performance.

To assess how transient high-frequency events contribute to stimulus decoding independent of electrode-level aggregation, each detected event was represented as a feature vector spanning morphology, temporal alignment, and state dimensions.

Decoding performance varied markedly across feature groups (**Figure 5A**). Morphology-related features, including amplitude, center frequency, frequency span, number of cycles, and duration, yielded decoding performance near chance (1/6), indicating that local transient-event structure carries relatively little category-specific information. In contrast, low-frequency phase (state) and latency (temporal) supported substantially higher decoding accuracy across subjects.

**Figure 5.**
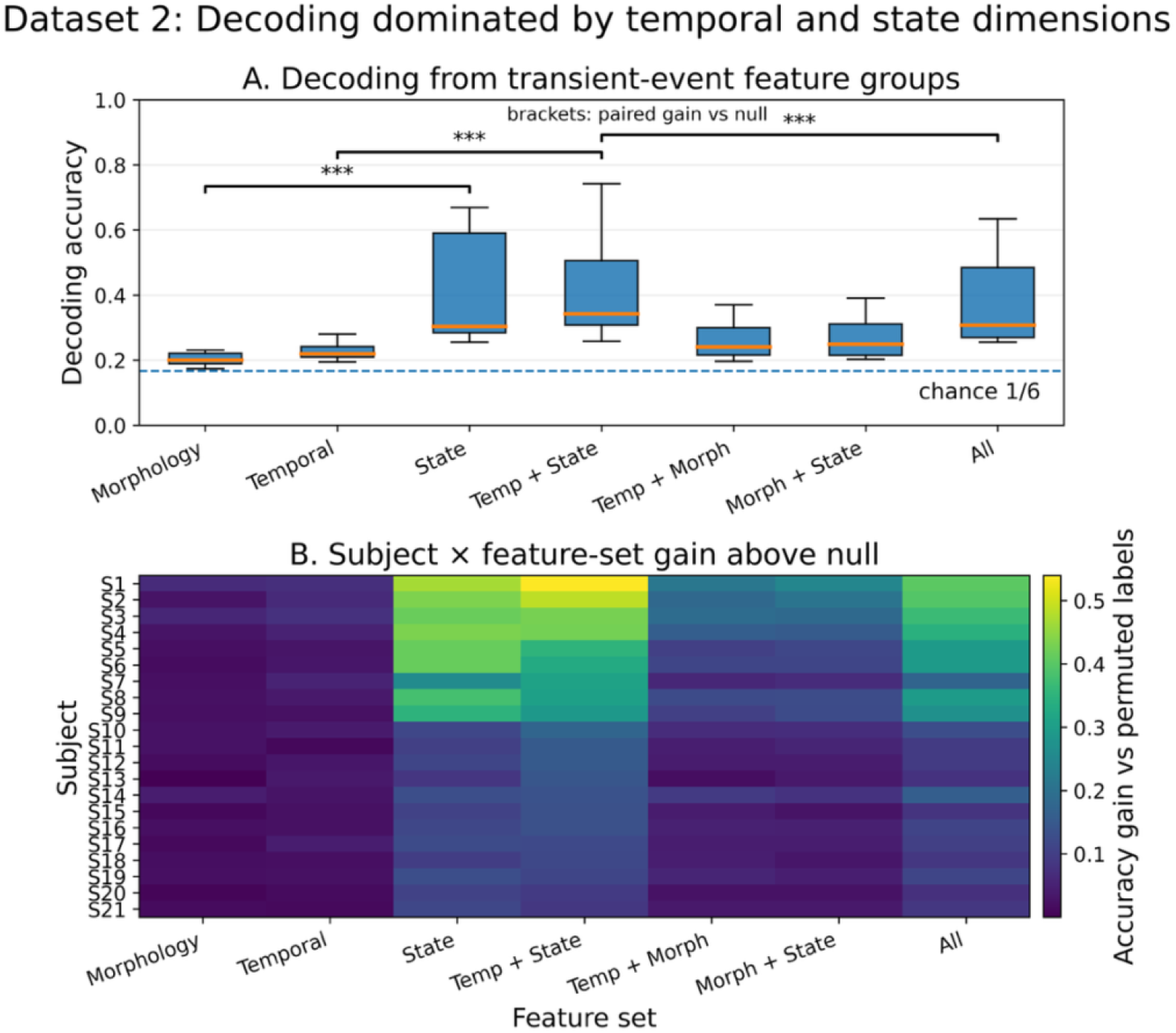
Decoding of transient high-frequency activity is dominated by temporal and state dimensions. **A**. Cross-validated decoding accuracy across subjects for models trained on grouped transient-event feature sets. Morphology-related features, including amplitude, center frequency, frequency span, number of cycles, and duration, yielded near-chance performance, whereas low-frequency phase (state) and latency (temporal) supported substantially higher decoding accuracy. Statistical annotations indicate paired comparisons of decoding gain relative to permuted-label null models (***, p < 0.001). **B**. Subject-by-feature-set heatmap of decoding gain relative to permuted-label null models. Temporal and state-related feature sets consistently produced larger gains across subjects, whereas morphology-related representations remained weak. Subjects are ordered by decoding gain in the temporal + state condition.

Combining temporal and state features produced the strongest compact representation, matching or exceeding the performance of the full feature set. In contrast, adding morphology-related features did not improve decoding performance, indicating that additional local event properties contribute little beyond temporal alignment and network-state information.

These effects were highly consistent across subjects (**Figure 5B**). Temporal and state-related feature sets showed robust increases in decoding gain relative to permuted-label null models, whereas morphology-related representations remained uniformly weak.

Targeted paired comparisons confirmed this structure. Morphology-related features contributed significantly less information than state features (p < 10^−5^, rank-biserial = −1.0). Temporal + state models also significantly outperformed temporal features alone (p < 10^−5^, rank-biserial = −1.0), demonstrating that temporal alignment is insufficient without the accompanying network-state context. Notably, the temporal + state representation outperformed the full feature set (p < 10^−4^, rank-biserial = 0.91), indicating that inclusion of additional morphology-related features does not enhance, and may instead dilute, the decodable signal.

Together, these results demonstrate that stimulus-related information in transient HFA is not evenly distributed across event features, but is concentrated primarily in temporal alignment and low-frequency state dimensions.

These findings refine the interpretation suggested by **Figure 2**. The reduced category specificity observed in high-response electrodes does not imply that weaker responses are inherently more informative. Instead, the present analyses indicate that stimulus-related information is concentrated in specific dimensions of the neural response, particularly temporal alignment and low-frequency phase context, whereas morphology-related properties contribute relatively little. Thus, the stereotyped response components dominating high-response electrodes appear to reflect broadly shared visual activation rather than category-specific structure.

The feature-group analyses in **Figure 5** identified temporal alignment and low-frequency phase as the dominant dimensions contributing to decoding performance, while morphology-related properties contributed relatively little. However, these analyses treated transient-event features primarily at the level of individual events and electrodes. This raised a critical question: does the observed contribution of low-frequency phase reflect a local property of individual transient events, or does informative structure emerge from the distributed organization of phase across cortical networks? To address this, we next constructed network-level representations based on the instantaneous low-frequency phase and amplitude measured simultaneously across electrodes at the time of each transient HFA event.

To isolate the contribution of distributed phase organization from distributed response strength, network-level phase representations were compared against matched amplitude-based representations extracted from the same transient-event timepoints and electrode populations. Unlike temporal descriptors such as event latency, which are defined relative to individual transient occurrences and therefore do not naturally yield a shared multielectrode representation across all events, instantaneous low-frequency phase and amplitude can both be sampled simultaneously across electrodes at each event timepoint. This enabled a direct comparison between distributed phase configuration and distributed response magnitude under equivalent network-level sampling conditions.

Decoding was then performed using grouped cross-validation in which all transient events originating from the same trial were assigned exclusively to either the training or test set, preventing trial-level leakage and ensuring that decoding reflected generalization across trials rather than repeated event structure.

Distributed low-frequency phase organization yielded substantially higher cross-trial decoding performance than matched amplitude-based network representations (**Figure 6A**). Using grouped cross-validation in which all transient events from a given trial were assigned exclusively to either training or test sets, low-frequency phase increased decoding gain above permutation null in 19 of 21 subjects (Wilcoxon signed-rank test, p < 0.001, rank-biserial correlation r = 0.86; **Figure 6A**).

**Figure 6.**
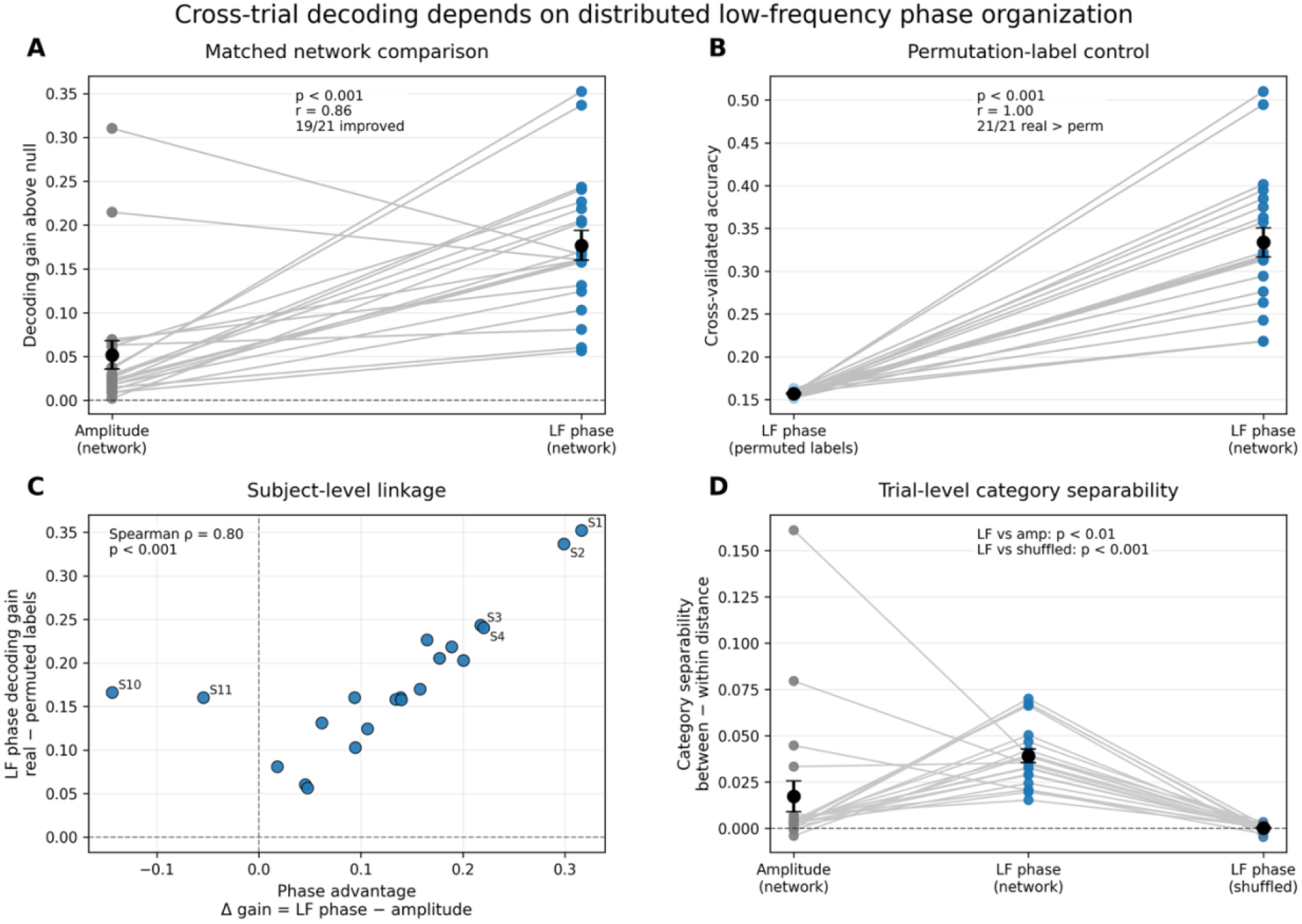
Cross-trial decoding depends on distributed low-frequency phase organization. **A**. Subject-level paired comparison of decoding gain above permutation null using grouped cross-validation, in which all transient events originating from the same trial were assigned exclusively to either training or test sets. Each line represents one subject. Distributed low-frequency phase (LF phase) representations yielded consistently higher decoding gain than matched amplitude-based network representations extracted from the same transient-event timepoints and electrode populations (Wilcoxon signed-rank test, p < 0.001, rank-biserial correlation r = 0.86; 19/21 subjects improved). Black markers indicate mean ± s.e.m. across subjects. **B**. Permutation-label control analysis. Decoding performance for distributed LF-phase network representations was compared against trial-wise label-permuted null estimates generated by randomly shuffling stimulus labels prior to classifier training. Real LF-phase representations consistently exceeded permutation-based decoding performance across all subjects (Wilcoxon signed-rank test, p < 0.001, rank-biserial correlation r = 1.00; 21/21 subjects showed higher real than permuted-label accuracy), indicating that decoding performance reflects stimulus-related structure rather than statistical bias or trial-count structure alone. **C**. Relationship between LF-phase decoding advantage and LF-phase decoding gain across subjects. Subjects exhibiting larger decoding gains for LF phase relative to amplitude also showed larger decoding gains above permutation-label null estimates (Spearman ρ = 0.81, p < 0.001), linking the relative advantage of LF-phase representations to overall decoding performance. **D**. Trial-level category separability measured as the difference between between-category and within-category distances in network feature space. Distributed LF-phase representations produced greater category separability than amplitude-based representations and phase-shuffled controls (LF phase vs amplitude: p < 0.01; LF phase vs shuffled: p < 0.001), indicating that low-frequency phase organization enhances the representational separation of visual categories across trials.

To verify that decoding performance reflected stimulus-related structure rather than statistical biases arising from feature dimensionality or trial organization, we compared LF-phase decoding against trial-wise permutation-label null estimates generated by randomly shuffling stimulus identities prior to classifier training. Real LF-phase representations consistently exceeded permutation-based decoding performance across all subjects (Wilcoxon signed-rank test, p < 0.001, rank-biserial correlation r = 1.00; **Figure 6B**), confirming that decoding performance was not attributable to chance structure in the dataset.

Across subjects, the magnitude of the LF-phase decoding advantage relative to amplitude was strongly associated with overall LF-phase decoding gain above permutation null (Spearman ρ = 0.81, p < 0.001; **Figure 6C**). Subjects exhibiting larger improvements from LF-phase representations relative to amplitude also showed larger decoding gains relative to trial-wise permutation-label baselines, linking the relative advantage of distributed phase organization to overall decoding performance.

To examine whether informative structure depended specifically on the distributed organization of phase across electrodes, we quantified category separability after independently shuffling phase values within each electrode while preserving marginal phase statistics and feature dimensionality. Distributed LF-phase representations produced significantly greater category separability than both amplitude-based representations and phase-shuffled controls (LF phase vs amplitude: p < 0.01; LF phase vs shuffled: p < 0.001; **Figure 6D**), indicating that low-frequency phase organization enhances the representational separation of visual categories across trials.

Together, these results demonstrate that stimulus-related information is not primarily determined by the local properties of individual transient HFA events, but instead depends on the distributed low-frequency network state in which those events occur. Notably, the decoding gains achieved by distributed phase representations exceeded those obtained from local transient-event features, suggesting that network-level phase organization captures structure not accessible at the level of individual electrodes.

## Discussion

Our findings indicate that stimulus-related information in transient HFA is not primarily reflected in local response magnitude or detailed event morphology, but instead emerges from distributed, state-dependent patterns across cortical networks. Across multiple levels of analysis, we found that individual transient-event features carried only modest information (**Figure 4**), whereas temporal alignment and low-frequency phase consistently contributed to decoding performance (**Figure 5**). Critically, decoding improved substantially when transient events were represented as distributed low-frequency phase configurations across electrodes (**Figure 6**), indicating that informative structure depends on coordinated network organization rather than isolated local event properties.

A key observation motivating this framework is that response magnitude alone does not capture category-specific structure (**Figure 2**). Electrodes with strong evoked responses exhibited highly stereotyped activity patterns across stimulus categories, whereas electrodes with weaker responses displayed more heterogeneous and temporally structured dynamics. This dissociation suggests that large-amplitude responses primarily reflect broadly shared components of visual processing, while stimulus-related structure resides in more subtle variations of the signal. Importantly, this does not imply that weaker responses are intrinsically more informative, but rather that informative structure is not captured by response magnitude alone.

By decomposing HFA into transient events and quantifying their temporal, spectral, and state-dependent properties, we were able to evaluate how different dimensions of neural activity contribute to stimulus-related information (**Figure 4A**). Individual transient-event features carried modest but reliable information about stimulus identity, consistent with a distributed representational scheme in which no single descriptor is sufficient for accurate classification (**Figure 4B**). Information varied substantially across subjects, indicating that anatomical sampling and electrode coverage strongly constrain the amount of observable stimulus-related structure (**Figure 4C**).

Within this feature space, not all dimensions contributed equally. Decoding performance was dominated by temporal alignment and low-frequency phase, whereas morphology-related properties, including frequency span, duration, and cycle structure, contributed relatively little (**Figure 5A**). Temporal features alone were insufficient for decoding, but improved performance when combined with low-frequency phase, suggesting that the timing of transient HFA events becomes informative primarily in relation to ongoing network state. Together, these findings indicate that stimulus-related information is not primarily reflected in detailed local event morphology, but in the temporal and state-dependent organization of transient activity.

At the network level, this effect became even more pronounced. Representing each transient event as a multielectrode low-frequency phase configuration yielded substantially higher decoding performance than matched amplitude-based representations extracted from the same event timepoints and electrode populations (**Figure 6A**). Importantly, this advantage depended on distributed cross-electrode phase structure rather than individual phase values alone, as disrupting phase relationships across electrodes abolished decoding performance (**Figure 6B**). These findings suggest that stimulus-related information in transient HFA is not localized within isolated event properties, but instead emerges from coordinated network dynamics distributed across cortical regions. Such coordination is consistent with the idea that low-frequency activity organizes neural activity across spatial scales, facilitating communication between distant cortical areas^13^.

This interpretation is consistent with prior work linking low-frequency activity, phase-amplitude coupling, and coordinated cortical dynamics to large-scale modulation of excitability and communication across neural populations^1,11,17,20–26^. and enable large-scale integration of distributed signals. In this context, low-frequency phase may provide a temporally structured network state within which transient HFA events occur, constraining the representational geometry of stimulus-related activity across trials. The near-simultaneous occurrence of transient HFA across distant cortical regions, despite conduction delays, further supports the possibility that these dynamics reflect coordinated interactions mediated through common inputs or distributed cortico-cortical and thalamocortical circuits. Aligning neural activity to specific phase relationships may enhance information transmission and facilitate large-scale integration of distributed signals^1,17^.

Importantly, the transient HFA events examined here should not be interpreted as definitive evidence of sustained oscillatory generators, nor can they be conclusively classified as classical high-frequency oscillations (HFOs). These events instead represent transient increases in high-frequency activity characterized according to their temporal, spectral, and state-dependent structure during visual processing. While some events may reflect genuine rhythmic synchronization or HFO-like activity, others may arise from asynchronous population dynamics or broadband spectral fluctuations. Recent work has emphasized that transient high-frequency events can emerge even in signals lacking stable oscillatory generators, and that event-detection approaches may capture structured but non-stationary neural transients rather than canonical oscillations alone^10^. Within this context, our framework is intentionally agnostic regarding the precise physiological origin of individual events. Rather than attempting to determine whether each event constitutes a true oscillation or HFO, we focus on whether their distributed temporal and phase-dependent organization carries stimulus-related information. The strong dependence of decoding performance on distributed low-frequency phase organization suggests that these transient HFA events, regardless of their precise mechanistic origin, are embedded within coordinated large-scale network dynamics.

The present findings therefore support a framework in which stimulus-related structure in transient HFA emerges through interactions between local cortical activity and distributed low-frequency network organization. In this framework, transient HFA events may reflect the activity of local cortical assemblies embedded within broader network states that constrain the temporal organization of neural activity across spatial scales. More broadly, these results illustrate how supervised learning and information-theoretic approaches can be used not only for prediction, but also for probing the representational structure of electrophysiological activity and evaluating the relative contribution of distinct neural dimensions to decoding performance. While these approaches do not establish causal mechanisms directly, they provide a complementary framework for generating mechanistic hypotheses regarding distributed neural coding and large-scale coordination. These observations may also have implications for understanding neural dynamics in both clinical and translational contexts.

Distributed phase-dependent organization of transient HFA may provide a useful framework for investigating cognitive processing in neurological and psychiatric conditions, including epilepsy, schizophrenia, and neurodegenerative disease and disorders affecting semantic processing^27–30^. Likewise, the strong decoding performance achieved using distributed low-frequency phase representations suggests potential applications for future brain-computer interface systems capable of decoding perceptual or cognitive states beyond traditional motor paradigms^31–37^. An important direction for future work will be the integration of these data-driven representations with biophysically grounded circuit models^38–42^. Detailed thalamocortical and cortical network models have demonstrated how interactions between neuronal populations, synaptic dynamics, and laminar organization give rise to coordinated oscillatory activity across multiple frequency bands^8,43–45^. Linking transient HFA dynamics and distributed phase organization to specific circuit mechanisms may therefore provide a pathway toward more mechanistic accounts of large-scale neural coordination and its disruption in neuropsychiatric disorders^30,46,47^.

In summary, our findings indicate that stimulus-related structure in transient HFA is not primarily determined by local response magnitude or detailed event morphology, but instead emerges most strongly through distributed, phase-dependent network organization shaped by anatomical sampling and ongoing cortical state. These results provide a framework for understanding how transient high-frequency neural activity contributes to sensory representations through coordinated dynamics distributed across space and time.

## Methods

### Participants and experimental procedures

#### Participants

21 patients monitored for pre-surgical evaluation of epileptic foci were included in the study (5 females, mean age 35±11). All participants gave fully informed consent, including consent to publish, according to NIH guidelines, as monitored by the institutional review board at The Feinstein Institutes for Medical Research, in accordance with the Declaration of Helsinki.

#### Data Acquisition

Recordings were conducted at North Shore University Hospital, Manhasset, NY, USA. Electrodes were either subdural grids/strips placed directly on the cortical surface and/or stereotactically implanted depth electrodes (Ad-Tech Medical Instrument, Racine, Wisconsin, and PMT Corporation, Chanhassen, Minnesota). Subdural contacts were 3 mm in diameter and 1 cm spaced (center to center distance), depth contacts were 2 or 1 mm in diameter and 2.5 or 5 mm spaced. Out of the 21 patients included in the study, 9 were implanted with both subdural grids/strips and depth electrodes and 12 were implanted only with depth electrodes (see Table S1 for individual specification). The iEEG signals were acquired at 512 Hz or 3 KHz using a clinical recording system (Xltek Quantum 256, Natus Medical) or a Tucker Davis Technologies (TDT) PZ5M module, referenced to a vertex screw or a subdermal electrode. Electrical pulses were sent upon stimuli onsets and recorded along with the iEEG data for precise alignment of task protocol to neural activity.

#### iEEG electrode localization

Prior to electrode implantation, patients were scanned with a T1-weighted 1 mm isometric structural magnetic resonance imaging (MRI) on a 3 Tesla scanner. Following the implant, a computed tomography (CT) and a T1-weighted anatomical MRI scan on a 1.5 Tesla were collected to enable electrode localization. Electrodes were localized and visualized using the iELVis toolbox^48^, which makes use of BioImage Suite, FSL, Freesurfer, and custom code^49,50^. Briefly, the electrodes were semi-automatically localized on the post-implantation CT, co-registered to the preoperative MRI scan using FSL’s BET and FLIRT algorithms^51^. The standard brain provided by Freesurfer (fsaverage) was used to project electrodes from all patients onto a single template. Colored labels of the cortical surface as presented in **Figure 1B** were derived from surface-based Glasser^52^ atlas.

#### Experimental Procedures

Participants were seated in bed in front of an LCD monitor. Stimuli were embedded in a 60 cm gray square and included color images from 6 categories: animals, patterns, people, places, tools, or words. Each category included 10 unique pictures, which were each presented 6 times. Images were flashed at a fixed pace of 1 Hz, each image was presented for 250 milliseconds and followed by a fixed inter stimulus interval of 750 milliseconds. To assure task engagement, participants were instructed to maintain fixation during the entire task and push a button if a picture repeated itself. The task included 360 trials.

### Signal preprocessing

#### iEEG Signal Preprocessing and HFA Estimation

We focused on the broad-band high-frequency activity signals (HFA, 70-150Hz) which have been demonstrated to relate to aggregate firing rate in human^53–55^ and non-human primates^56,57^. This signal is also termed high-gamma^58^.

Signals that were initially recorded at a sampling rate of 512 Hz, 1kHz, or 3kHz were down-sampled to 500 Hz for consistency. Raw time-courses and power spectra of all channels were initially manually inspected for noticeable abnormal signals and other contaminations, and channels appearing as highly irregular were excluded from further analysis. In addition, channels identified as located over the seizure onset zone by an epileptologist were excluded.

To extract HFA power modulations, we segmented the data within a 4 second window relative to image presentation (-2 sec. - + 2 sec.) and convolved the signals with 6-cycle Morlet wavelets. We calculated the power as the squared absolute value of the resulting convolved signals for each electrode and averaged the power across the HFA frequency band (70 - 150 Hz). To detect electrodes that showed an increase in HFA power following the image presentation, we averaged the power over a 500 millisecond time window following the image presentation and calculated the t-test between the averaged signal and baseline power value (250 milliseconds prior to image presentation). All data preprocessing and analyses were carried out using customized Matlab and Python codes.

#### Detection of Transient Oscillation Events

We extracted spectral events using 6-cycle Morlet wavelets for the entire data using the BOSC method^19^. The 6-cycle Morlet wavelets were chosen to provide an adequate compromise between time and frequency resolution. We used linearly spaced frequencies with 1 Hz frequency increments, ranging from 70-150 Hz, to compute the wavelets. The power time-series of each wavelet transform was normalized by median power across the full recording duration.

All local peaks were assessed to determine whether their power value exceeded a threshold. We used a moderate power threshold (4× median of each individual frequency) to determine the occurrence of moderate-to-high-power events. We only considered HFA events occurring within a window of 0-0.5 seconds following the image presentation for further analysis and feature extraction.

Instantaneous phase was calculated by filtering the entire recording via convolution with a complex Morlet wavelet of width 6 and extracting the instantaneous phase at the appropriate frequencies (f-phase = delta - 0.5-4 Hz; theta - 4-8 Hz; alpha - 8-12 Hz; beta - 14-25 Hz). Then, the instantaneous phase values were extracted based on the timing of the HFA events.

Instantaneous phase was calculated by filtering the entire recording via convolution with a complex Morlet wavelet of width 6 and extracting the instantaneous phase at the appropriate frequencies (f-phase = delta - 0.5-4 Hz; theta - 4-8 Hz; alpha - 8-12 Hz; beta - 14-25 Hz). Then, the instantaneous phase values were extracted based on the timing of the HFA events.

### Dataset Construction

Three complementary feature representations were constructed to evaluate stimulus-related information at different organizational scales.

Dataset 1 captured feature values independently across trials and electrodes. For each subject and feature type, matrices were constructed with trials as rows and electrodes as columns, allowing assessment of the information carried by individual transient-event descriptors.

Dataset 2 represented each detected transient HFA event as a multidimensional feature vector spanning morphology, temporal alignment, and state-related properties. Features were grouped into morphology (center frequency, frequency span, duration, cycle count, amplitude), temporal (latency), and state (low-frequency phase) categories for grouped decoding analyses.

Dataset 3 captured distributed network-level representations by measuring instantaneous low-frequency phase or amplitude simultaneously across electrodes at the time of each transient HFA event.

### Feature Extraction and Grouping

For each detected HFA event, we extracted features describing event morphology (amplitude, center frequency, frequency span, duration, number of cycles), timing (latency relative to stimulus onset), and state (instantaneous low-frequency phase at event onset in delta, theta, alpha, and beta bands). Features were computed on a single-trial basis for each electrode and grouped into morphology, timing, and state categories for subsequent analyses.

### Decoding Analysis

Classification analyses were performed using Random Forest classifiers implemented in scikit-learn^58^. Decoding performance was evaluated independently for each subject using repeated stratified cross-validation. For Dataset 1 and Dataset 2 analyses, decoding was performed using stratified train-test splits repeated across multiple cross-validation runs, and performance estimates were averaged within subject.

For Dataset 3 analyses, grouped cross-validation was used such that all transient HFA events originating from the same trial were assigned exclusively to either the training or test set, preventing trial-level leakage across folds. Decoding performance was quantified as classification accuracy on held-out data.

### Null Model and Decoding Gain

Null distributions were generated independently within subject by randomly permuting stimulus labels and repeating the full decoding procedure using identical cross-validation parameters. Decoding performance was quantified as decoding gain (Accuracy_true - Accuracy_null_mean), controlling for biases related to class structure, feature dimensionality, and classifier variance.

### Statistical Testing

All statistical analyses were performed at the subject level. For each feature or feature group, decoding gain was computed per subject and tested against permutation-corrected null performance using two-sided Wilcoxon signed-rank tests. Pairwise comparisons between feature groups were performed using the same test. P-values were corrected for multiple comparisons using the Holm-Bonferroni method. Effect sizes were quantified using matched-pairs rank-biserial correlation.

### Control Analyses

Control analyses verified that decoding performance collapsed toward chance under label permutation. For network-level analyses, additional controls disrupting distributed phase configuration while preserving marginal phase statistics and feature dimensionality produced marked reductions in decoding performance, confirming that decoding depended on distributed phase organization rather than electrode-wise phase values alone.

### Data Visualization and Reporting

Results were visualized using subject-level distributions and mean decoding gain across conditions. Statistical heatmaps were represented as −log_10_(p), where 1.3 corresponds to p = 0.05.

